# Capsule protects against intracellular killing and enables vascular endothelial cell translocation during invasive pneumococcal disease

**DOI:** 10.1101/2021.02.10.430484

**Authors:** Terry Brissac, Eriel Martínez, Katherine L. Kruckow, Ashleigh N. Riegler, Feroze Ganaie, Hansol Im, Sayan Bakshi, Nicole M. Arroyo-Diaz, Brady L. Spencer, Jamil S. Saad, Moon H. Nahm, Carlos J. Orihuela

## Abstract

*Streptococcus pneumoniae* (*Spn*) is a leading cause of invasive disease. Chief among its virulence determinants is capsular polysaccharide which protects the bacterium from phagocytosis. While 100 antigenically distinct capsule types are produced by *Spn*, i.e. serotypes, only 20-30 are commonly associated with invasive disease. A frequency that suggests serotype-specific properties of the capsule influence virulence. Herein, we show capsule has strong antioxidant properties. Moreover, that this property promotes invasive disease by protecting *Spn* taken up by vascular endothelial cells during bacteremia from endosome-killing and enhancing the translocation rate into organs. Crucially, isogenic capsule-switch mutants of *Spn* varied considerably in their resistance to H_2_O_2_-killing in culture and measured levels correlated positively with intracellular survival rates *in vitro*, organ invasion rates *in vivo*, and epidemiologically-established human attack rates for the corresponding serotype. The amount of capsule produced and specific biochemical features of a serotype, such as acetylation, also influenced *Spn* resistance to oxidative stress. Autolysin-mediated shedding was also found to be necessary, indicating that capsule worked as a distal sink for reactive oxygen species. Our results outline a new role for capsular polysaccharide, as an intracellular antioxidant. They help to explain why certain serotypes of *Spn* have greater propensity for human disease.

## INTRODUCTION

*Streptococcus pneumoniae* (*Spn*) is the leading cause of community-acquired pneumonia. As result, it is also a leading cause of invasive diseases including bacteremia and meningitis. Young infants, the elderly, immunocompromised individuals, and those who recently experienced a viral respiratory tract infection, are particularly susceptible to the severe respiratory tract infections that can lead to invasive pneumococcal disease (IPD) (1, 2). Critically, mortality rates for the elderly with hospital-admitted pneumococcal pneumonia with bacteremia can be as high as 60% (3, 4). Thus, pneumococcal infection is a major medical problem. It is noteworthy that not all *Spn* are equally capable of causing IPD. To date, 100 biochemically and serologically distinct capsule types of *Spn* have been identified (5), of which only 25-30 are commonly associated with human disease (6, 7). Thus, the most obvious determinant of disease propensity is the capsule type carried by the infecting strain.

Polysaccharide capsule is a primary virulence determinant of *Spn* and numerous other pathogens (8). Non-encapsulated *Spn* can cause disease, but this is almost never life-threatening and generally restricted to the upper respiratory tract or the eye (9). The reason for this is that capsule protects the bacterium from host clearance *via* inhibition of complement deposition and by obscuring bacteria surface-attached host defense factors from their cognate receptors on immune cells (e.g. Fc portion of antibody), thereby blocking opsonophagocytosis (10, 11). In addition, the vast majority of *Spn* capsule types are negatively charged and this electrostatically repels immune cells (12, 13). Importantly, and for many of the same reasons, the capsule also impairs bacterial interactions with non-immune cells (14–16); an event that is critical for *Spn* establishment in the nasopharynx during colonization and progression of the disease (17). Pneumococci adjust for this inhibitor effect by modulating the amount of capsule produced via phase-variation (18), shedding their capsule when in close contact with epithelial cells (16), the latter due to exposure to cationic peptides released from these cells (19), and the incorporation of stalk-like elements within surface proteins to allow for the extension of effector domains beyond the capsule layer (20). Critically, pneumococci within the bloodstream are always encapsulated, as without capsule they are quickly cleared by immune cells. Thus, capsule is present when pneumococci in the bloodstream interact with vascular endothelial cells, an event preceding organ invasion.

*Spn* is the prototypical extracellular pathogen. Yet, we also know that *Spn* can be taken up by non-immune cells such as mucosal epithelial cells and vascular endothelial cells (VEC) (21). Whereas the latter has long been appreciated to be an essential step towards *Spn* translocation across the blood-brain barrier and the pathogenesis of meningitis (22), VEC translocation by *Spn* is now appreciated as having a vital role in other aspects of disseminated organ damage during bacteremia such as cardiac damage (23). Here, and in stark contrast with the general notion that the capsule is inhibitory of *Spn* invasion processes, we demonstrate that capsular polysaccharide is required for efficient *Spn* translocation across VEC. We go on to identify a new role for this classic virulence determinant, as an intracellular antioxidant, and show this property subsequently promotes *Spn* survival within VEC, enhances the rate of translocation across cell barriers, and helps to explain the propensity for disseminated organ invasion by distinct serotypes. Importantly, we extend our findings to other encapsulated pathogens and this expands our understanding of bacterial pathogenesis in a larger context.

## RESULTS

### Capsule mediates *Spn* transmigration through a vascular endothelial cell layer

Capsule has been shown to starkly reduce *Spn* interactions with VEC (14, 16, 24), yet it does not completely abolish *Spn* invasion. How capsule impacts *Spn* fate once internalized was unknown. We developed an *in vitro* Transwell^®^ assay using mouse cardiovascular endothelial cells (MCEC) that allowed for step-wise analyses of the bacterial transmigration process (Fig. 1A). MCEC were chosen as the prototype host cell since they formed confluent leak-free monolayers (Fig. S1), and during bacteremia *Spn* must cross this cell type to invade the myocardium to cause cardiac damage (23).

**Figure 1.**
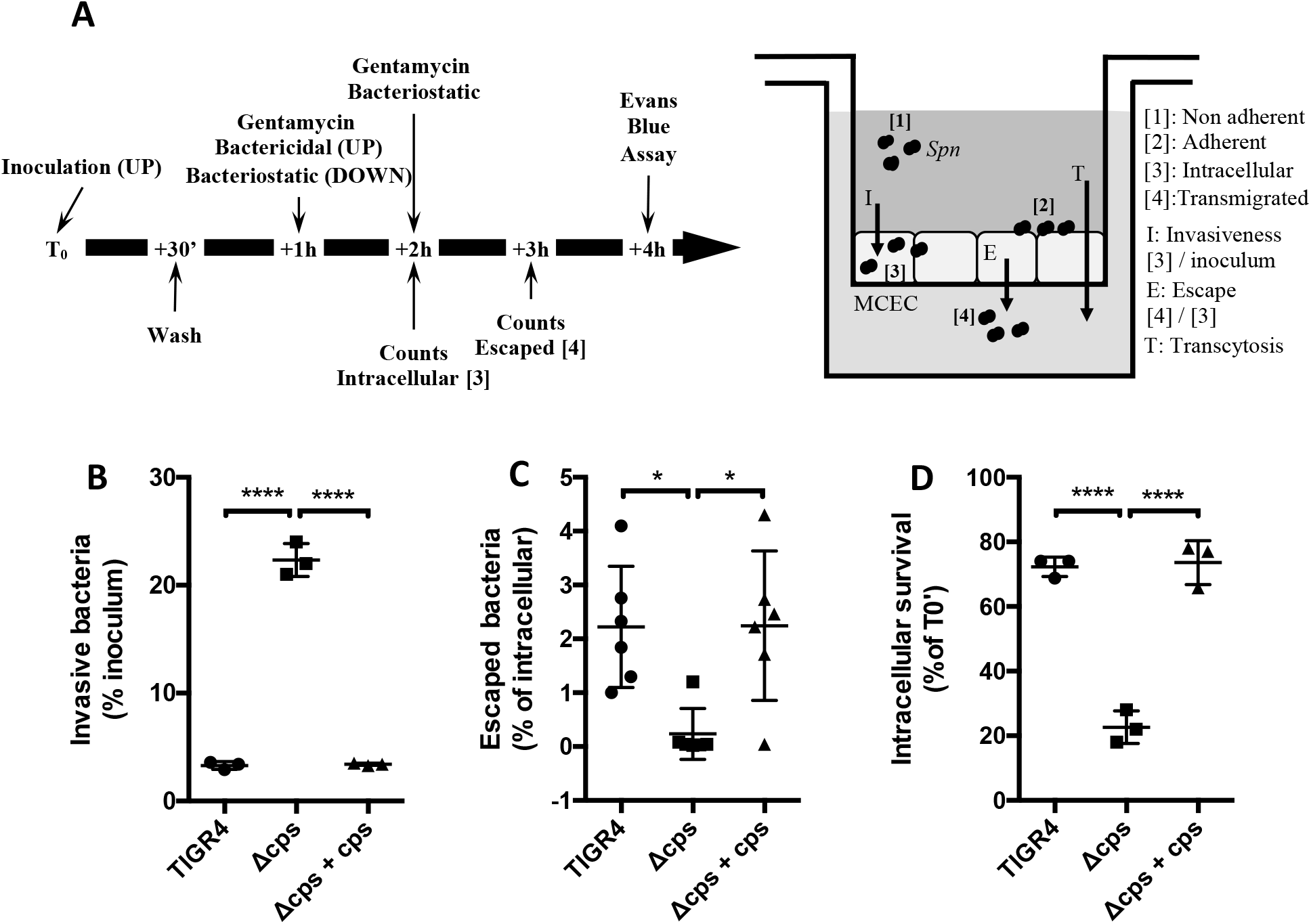
Capsule increases intracellular survival and bacterial escape from vascular endothelial cells. **(A)** Experimental flow-chart and Transwell^®^ system set-up (MCEC, mouse cardiovascular endothelial cells). **(B-C)** Encapsulated *Spn* (TIGR4) are less internalized but escape at higher rate than non-encapsulated mutant (Δcps). **(D)** Capsule promotes *Spn* intracellular survival. Intracellular survival of designated bacterial strains was determined by normalizing CFU at T_0_+2h of treatment by CFU at T_0_+1h of treatment (T_0_’). Statistical analyses: Mann-Whitney U-test (B, C, D). Errors bars represent standard error of the mean.

Consistent with existing literature, an isogenic capsule deletion mutant of TIGR4 (Δ*cps*) was internalized by MCEC at considerably greater rates, >5-fold, than wild-type (WT) TIGR4 or the *cps* complemented strain (Δ*cps* + *cps*) (Fig. 1B). Yet, the percentage of WT TIGR4 or Δ*cps* + *cps* that successfully escaped from within cells to the basolateral surface of the monolayer was >10-fold greater than Δ*cps* (Fig 1C). We hypothesized that capsule enhanced VEC escape by prolonging bacterial intracellular survival. In support of this notion, gentamicin protection assays (25) showed that after 2 hours of treatment, killing specifically extracellular bacteria, the number of viable *Spn* recoverable from within MCEC was >3-fold higher for WT TIGR4 versus Δ*cps* (Fig. 1D).

### Capsule increased intracellular survival by increasing tolerance to oxidative stress

Following clathrin-mediated endocytosis, bacteria within phagosomes are exposed to multiple stressors meant to kill and degrade cargo; this includes reactive oxygen species (ROS) (26). Using bacterial culture media supplemented with 10 mM H_2_O_2_ (27); we observed that the presence of capsule conferred up to 30 minute delay in *Spn* killing (Fig. 2A, Fig. S2). To investigate whether the amount of polysaccharide present on the pneumococcal surface affected resistance to ROS, we constructed a mutant harboring a constitutive promotor (Pcat) upstream of the capsule operon (Pcat-*cps*) that was comparably weaker than the native version found in WT TIGR4 (28). FITC-dextran exclusion assay confirmed that Pcat-*cps* produced about half the amount of capsule produced by WT TIGR4 (Fig. S3). The reduced amount of capsule produced by Pcat-*cps* was enough to protect intracellular *Spn* when compared to Δ*cps*, despite being more susceptible to killing than WT TIGR4 (Fig. 2B). Additionally, and *in vitro*, the protection conferred by the Pcat-*cps* mutant when treated with H_2_O_2_ was significantly less than WT TIGR4 (Fig. 2C). We subsequently hypothesized that encapsulated bacteria were less susceptible to ROS through a buffering mechanism whereby oxygen-derived free radicals preferentially attacked the polysaccharide fibers. In support of this hypothesis, we observed that addition of purified serotype 4 capsule under physiological concentrations protected Δ*cps in vitro* (Fig. 2D). To confirm that the capsule was acting as a free radical scavenger, we measured the antioxidant ability of purified capsule using a nitroblue tetrazolium (NBT) reduction assay. The addition of capsular polysaccharide affected the kinetic of NBT reduction in a dose-dependent manner (Fig. 2E) supporting the notion that serotype 4 capsule had antioxidant properties. Subsequently, one- and two-dimensional nuclear magnetic resonance (NMR) data of purified type 4 capsular polysaccharide exposed to 10mM H_2_O_2_ for 30 minutes (Supplemental Fig S4) and 3 hr (Fig. 2F) showed that exposure to H_2_O_2_ resulted in specific structural/conformational changes as would be expected if capsule was indeed an antioxidant. These changes occurred at more than one site suggesting it was more than one specific biochemical moiety that was responsible for this effect.

**Figure 2.**
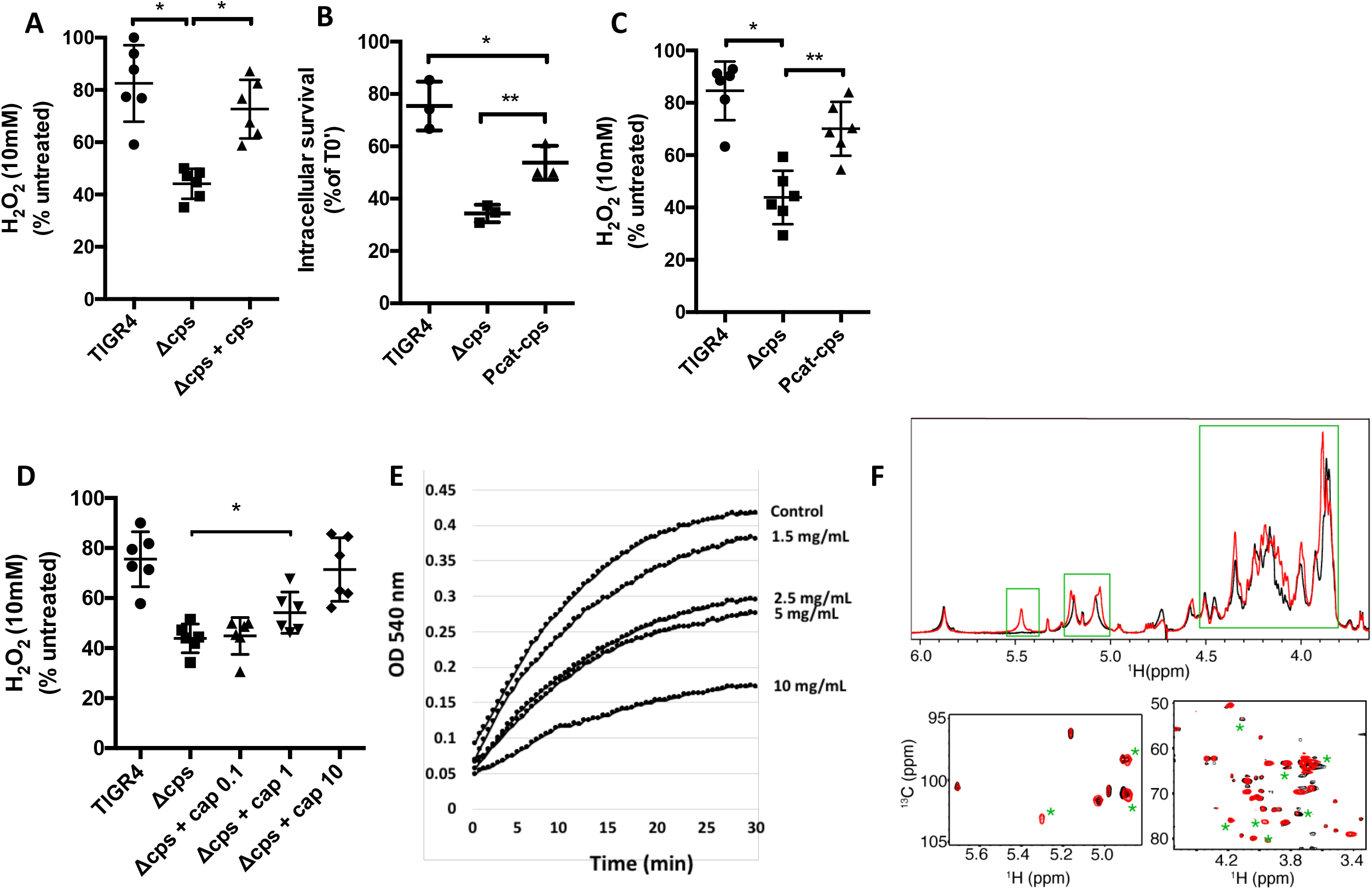
Capsule increased tolerance to oxidative stress. **(A)** Capsule presence increases tolerance to H_2_O_2_. Tolerance was determined by calculating the number of live *Spn* in THY supplemented with 10mM H_2_O_2_ compared to *Spn* incubated in plain THY after 15 minutes of incubation. **(B-C)** Capsule amount influences *Spn* intracellular survival and tolerance to H_2_O_2_. **(D)** Exogenous polysaccharide at 1mg/ml or higher protected un-encapsulated TIGR4 (Δcps) from killing by H_2_O_2_. **(E)** Purified capsule showed antioxidant properties in a NBT reduction assay. **(F)** ^1^H NMR (upper) and 2D ^1^H-^13^C HMQC spectra (lower) for untreated (black) and H_2_O_2_-treated (red) samples of serotype 4 PS at 50 °C. Peaks marked with green boxes or asterisks denote significant spectral changes. Statistical analyses: Mann-Whitney U-test (A-C), One way ANOVA with Tukey’s multiple comparison (D). Errors bars represent standard error of the mean.

### Biochemical properties of a serotype influence intracellular survival

At least 100 serologically distinct serotypes of *Spn* are known to exist. Strikingly, the escape rate of a panel of *Spn* clinical isolates positively correlated with the published attack rates, i.e. invasive disease per colonization event, for their corresponding capsule type (Fig. 3A). A finding which suggested that the impact of the capsule on the *Spn* translocation process strongly influenced disease. To dissect the contribution of capsule from the confounding effects of disparate genomes, we created isogenic capsule switch mutants of nine clinically relevant serotypes in the genetic background of TIGR4. As expected, we observed considerable variability in the capability of these capsule switch mutants to survive within MCEC cells (Fig 3B) and their resistance to 10 mM H_2_O_2_ *in vitro* (Fig 3C). Yet we also observed a very strong positive correlation between intracellular survival and H_2_O_2_ resistance of tested serotypes (Fig. 3D). To further establish the relevance of serotype-mediated resistance to oxidative stress on intracellular protection and bacterial translocation we took advantage of the difference observed in the intracellular survival of TIGR4 expressing its own type 4 capsule (TIGR4^ISO4^) and TIGR4 producing type 2 capsule (TIGR4^ISO2^). We found that TIGR4^ISO4^ had double the MCEC escape frequency of TIGR4^ISO2^ (Fig. 3D). The addition of exogenous purified capsule 4 to the media also protected Δ*cps* more efficiently than purified capsule 2 following exposure to 10mM of H_2_O_2_ (Fig. 3F). Finally, and as expected, capsule type 4 displayed greater antioxidant activity than capsule type 2 in our NBT reduction assay (Fig. 3G).

**Figure 3.**
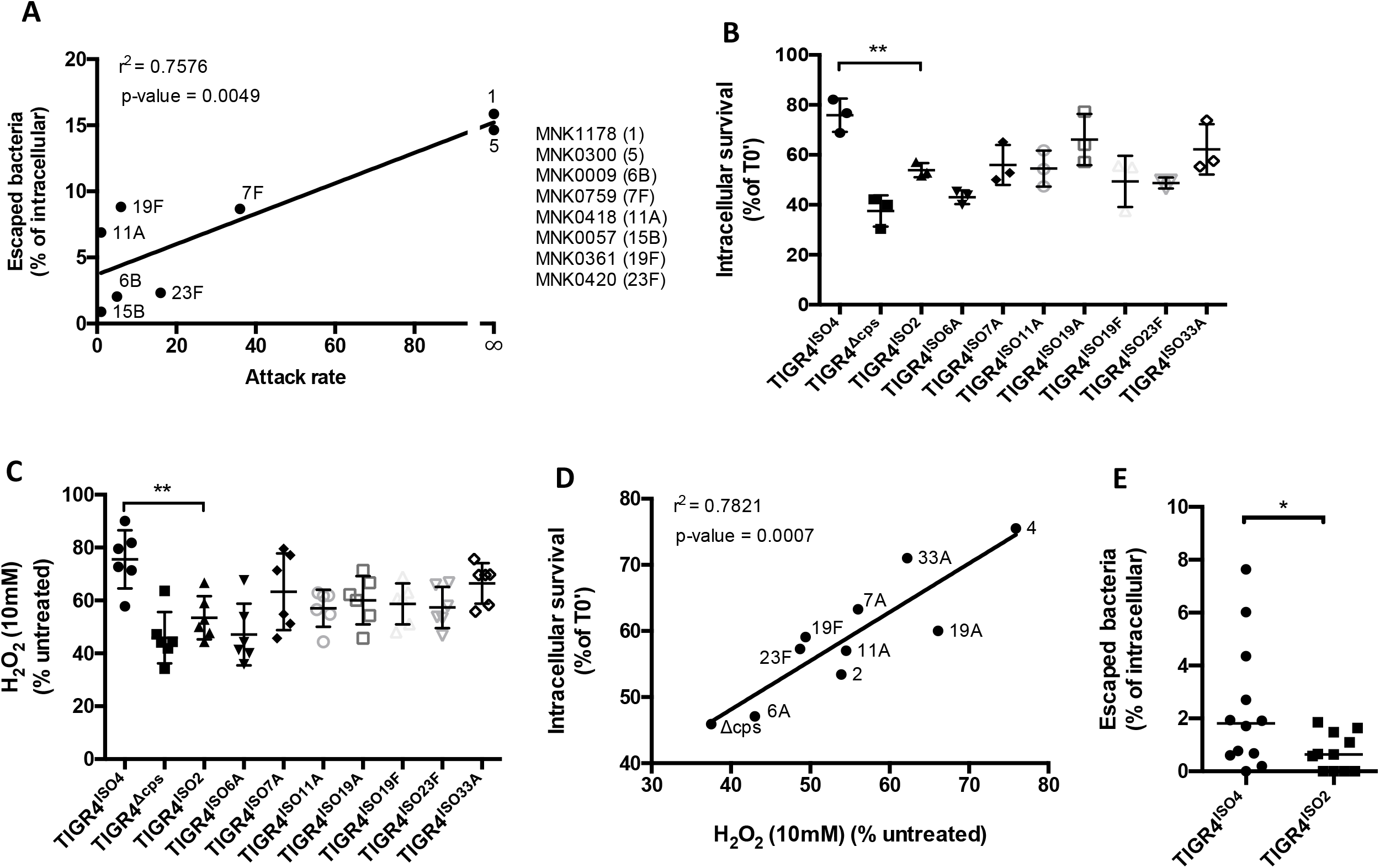
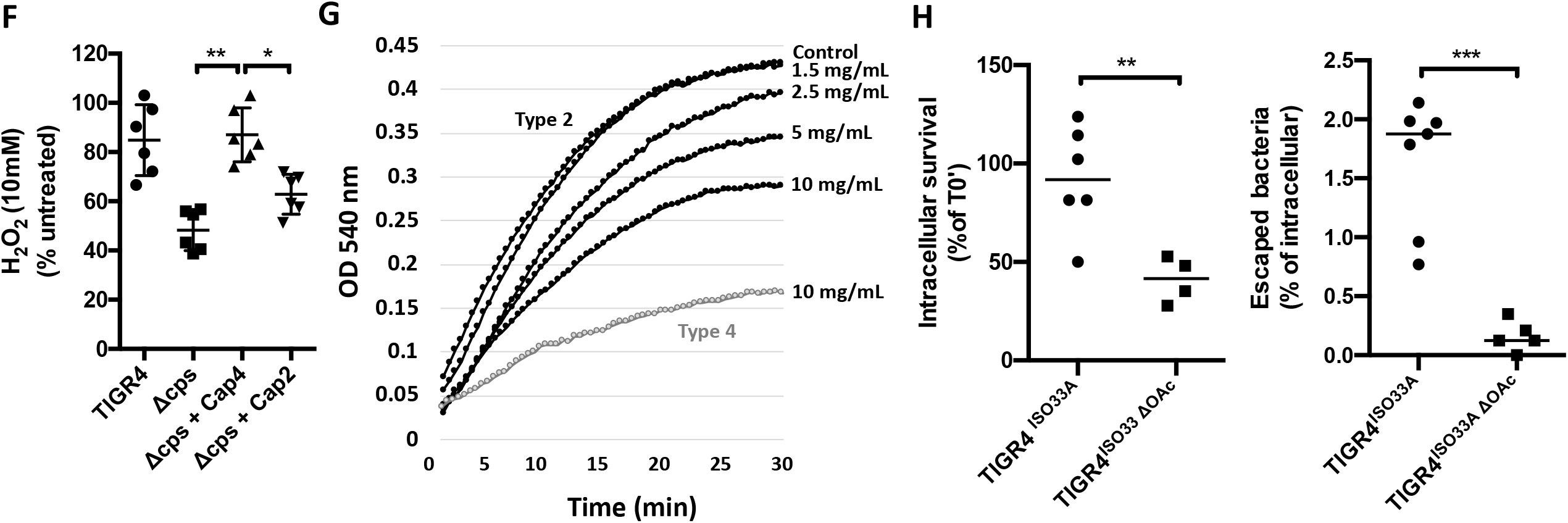
Capsule serotype influences oxidative stress tolerance. **(A)** Escape rate determined for a panel of clinical isolates selected among various serotype correlated with the attack rate reported for the corresponding serotype **(B-C)** TIGR4 expressing different capsule types showed variable intracellular survival and H_2_O_2_ tolerance. **(D)** Serotype intracellular survival correlates with oxidative stress tolerance. **(E)** TIGR4 expressing serotype 2 showed reduced escape rate compared with TIGR4 expressing its own serotype 4 capsule. **(F)** Exogenous capsule 2 showed less level of Δcps protection compared to exogenous capsule 4. **(G)** Capsule 2 showed less antioxidant capability compared to capsule 4 (see figure 3F) in a a NBT reduction assay. Capsule 4 10mg/mL (grey line) was plotted as a reference **(H)** TIGR4 expressing capsule O33A capsule showed higher intracellular survival and escape than the same strain expressing the O33 un-acetylated variant O33A ΔO-Ac. Statistical analyses: Spearman correlation (A, D) One way ANOVA with Tukey’s multiple comparison (B, C), Student’s *t*-test (E, H) Mann-Whitney U-test (F). Errors bars represent standard error of the mean.

One major difference between type 2 and type 4 capsule is the presence of N-acetylated groups on type 4 (12). To discern the effect of acetylation on the capability of the capsule to protect intracellular *Spn* we used TIGR4 strains expressing capsule type 33A (TIGR4^ISO33A^) and a mutant lacking the required capsule acetyltransferases (TIGR4^ISO33A ΔOAc^) (29). TIGR4^ISO33AΔOAc^ was starkly impaired for survival within MCEC cells when compared to TIGR4^ISO33A^ (Fig. 3H). Thus, the biochemical traits of each capsule type are a key factor in the ability of *Spn* to escape from within cells and thereby translocate across VEC to cause IPD.

### Capsule shedding promotes intracellular survival and bacterial escape

LytA is the major pneumococcal autolysin. It is the cell wall amidase responsible for *Spn* bile solubility and autolysis following exposure to cell wall acting antimicrobials (30). Importantly, LytA is also activated in the presence of antimicrobial peptides causing the release of cell wall-attached capsule without concomitant bacterial lysis. This non-lytic activity of LytA is called capsule shedding (19). Notably, a LytA deficient isogenic mutant of TIGR4 (Δ*lytA*) had reduced intracellular survival within MCEC (Fig. 4A) and had diminished escape (Fig 4B) rates. A double unencapsulated Δ*lytA* mutant (i.e. Δ*cps* Δ*lytA*) was not further attenuated for these traits (Fig. 4A and 4B), indicating that LytA-mediated shedding of the capsule and no other collateral effect of *lytA* deletion was required for the protective effect. These results suggested the free PSs released by LytA serve to neutralize ROS at distance. In agreement with this hypothesis Δ*lytA* showed a slightly but significant reduced resistance to H_2_O_2_ *in vitro* (Fig. 4C).

**Figure 4.**
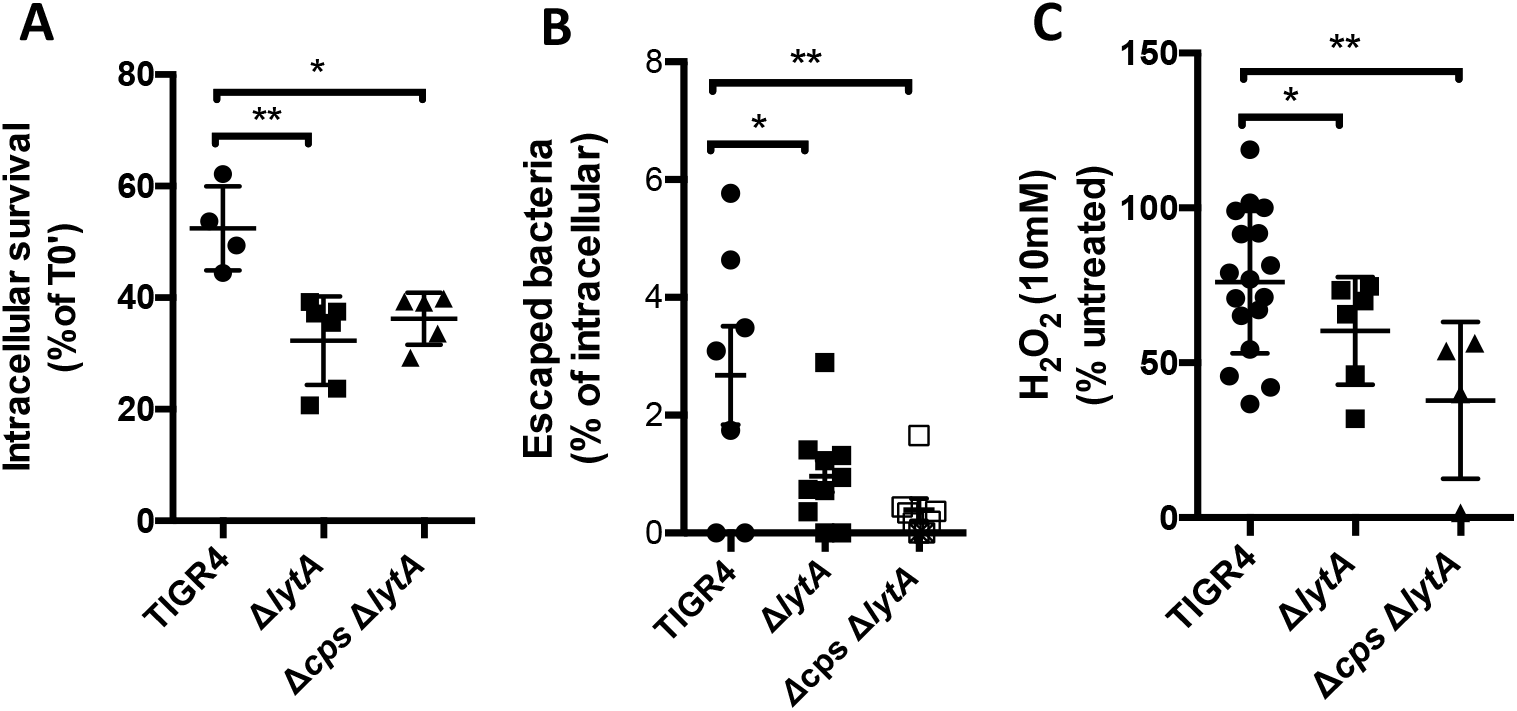
Capsule shedding promotes *Spn* transcytosis. **(A-B)** A shedding deficient mutant of TIGR4 (Δ*lytA*) showed reduced intracellular survival and escape capabilities compared to TIGR4 WT. **(C)** Δ*lytA* showed reduced tolerance to H_2_O_2_. Statistical analyses: Mann-Whitney U-test. Errors bars represent standard error of the mean.

### Capsule impacts disease presentation

Our results thus far suggest capsule-type conferred resistance to H_2_O_2_ would have a direct effect on organ invasion during disseminated infection. To test this possibility, we intraperitoneally (i.p.) challenged mice with TIGR4^ISO4^ or TIGR4^ISO2^ and examined the ability of each strain to translocate into the myocardium. This model was chosen as bacteria in the peritoneum continuously enter the circulation via the lymphatic system, avoiding the bottleneck that occurs in the spleen following intravenous injection, as reported by Ercoli et al (31). Moreover, invasion of the myocardium requires vascular endothelial cell translocation from the bloodstream (23). Following challenge, mice infected with TIGR^ISO4^ had bloodstream titers 100-fold lower than those challenged with TIGR^ISO2^ (Fig 5A), yet equivalent numbers of TIGR4^ISO4^ and TIGR4^ISO2^ were recoverable from perfused hearts of these mice. When normalized against the number of bacteria in the blood, TIGR4^ISO4^ was ~50-fold more efficient at invading the heart compared to TIGR4^ISO2^ (Fig. 5B). To rule out the possible positive effects of the matched genetic background and serotype for TIGR4^ISO4^ we also generated and tested isogenic mutants in a serotype 3 genetic background. WU2 expressing capsule types 4 and 2 (WU2^ISO4^ and WU2^ISO2^, respectively) showed similar results to the TIGR4 genetic background (Fig. 5C and 5D). Finally, and to ascertain if our laboratory results could help explain real-world disease rates, we tested for a correlation between observed capsule-mediated resistance to oxidative stress and published epidemiological attack rates for distinct serotypes of *Spn* as well as between observed intracellular survival rates and attack rates. Strikingly, we observed strong correlations between the two suggesting this was an important feature that determined pneumococcal propensity for invasive disease (Fig 6).

**Figure 5.**
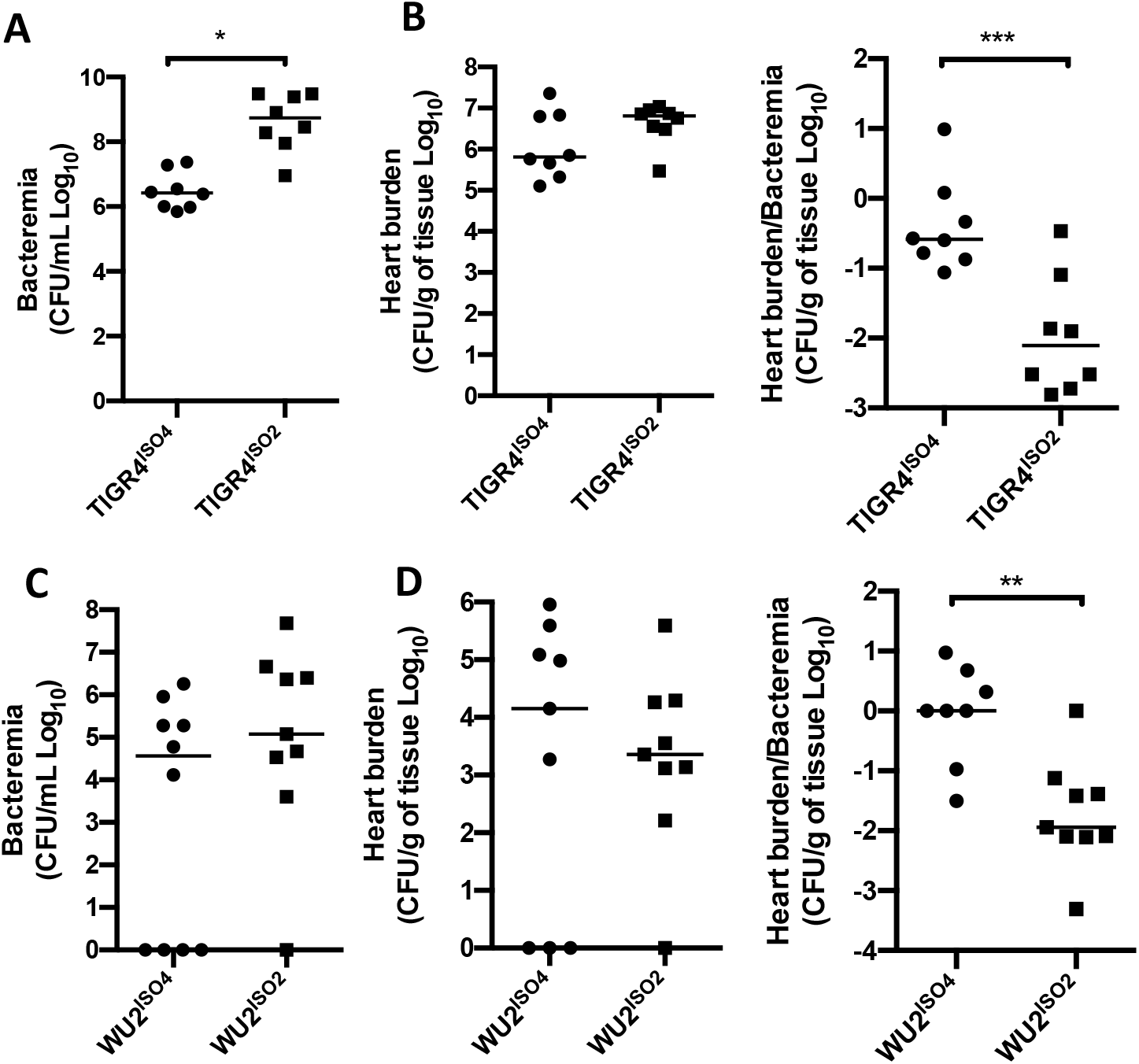
Capsule impact disease presentation. **(A, C)** C57B6 mice were challenged intraperitoneally using 1.0×10^5^ CFU/ml of TIGR4 (A) or WU2 (C) background isogenic capsule switches and bacteremia and heart titer were determined at time of sacrifice (T0+36h~42h). **(B, D)** Tissue invasion ability of strains was determined by normalization of heart titer by the bacteremia N=8. Statistical analyses: Student’s *t*-test. Errors bars represent standard error of the mean.

**Figure 6.**
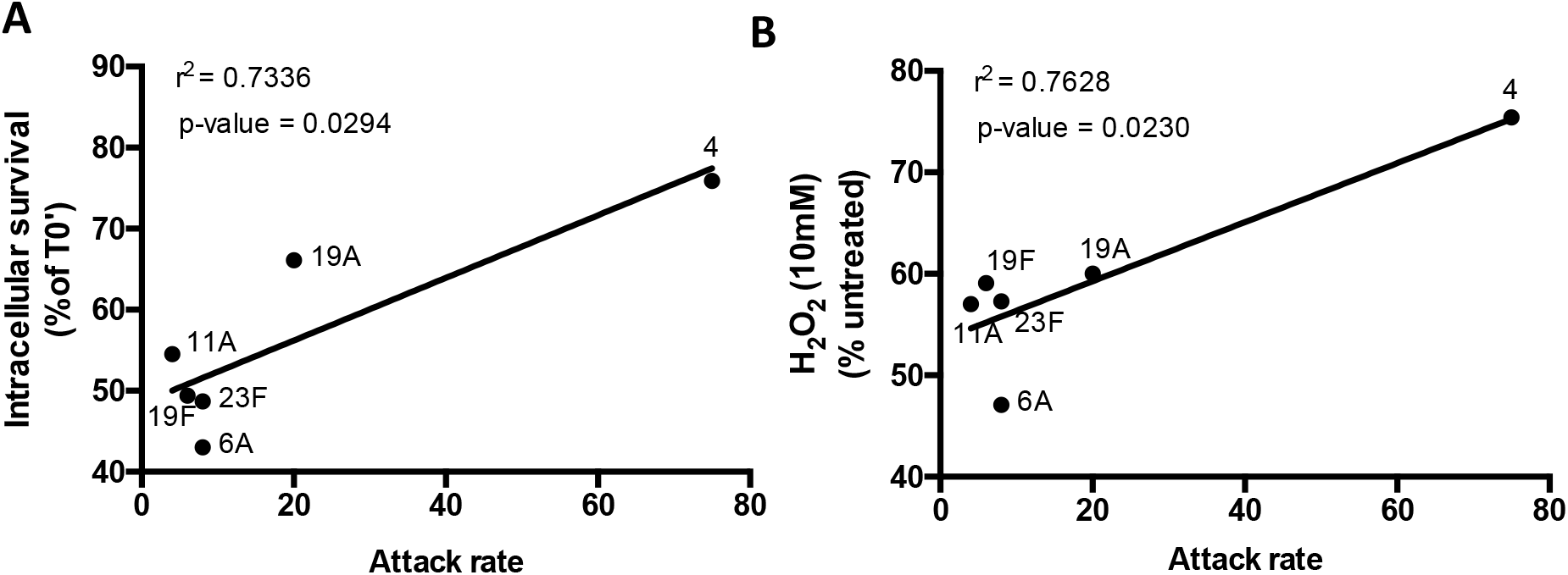
Intracellular survival and oxidative stress resistance of the capsule switch mutants correlate with the attack rate of clinical isolate serotypes. **(A-B)** Intracellular survival fate and hydrogen peroxide resistance of capsule switch mutants were plotted with the attack rates reported for each serotype. Statistical analyses: Spearman correlation.

### Capsule promotes transcytosis of other invasive pathogens

The production of a polysaccharide capsule is a common feature among many different bacteria capable of causing invasive disease. To test if our observations extended to other pathogens, we explored the role of the capsule on intracellular protection and translocation of *Streptococcus pyogenes* and *Staphylococcus aureus*. Notably, both of these pathogens escaped at higher rates than their respective non-encapsulated isogenic mutants, despite being initially outnumbered within the cells (Fig. 7). These results suggest our observations for *Spn* capsule are most likely broadly applicable.

**Figure 7.**
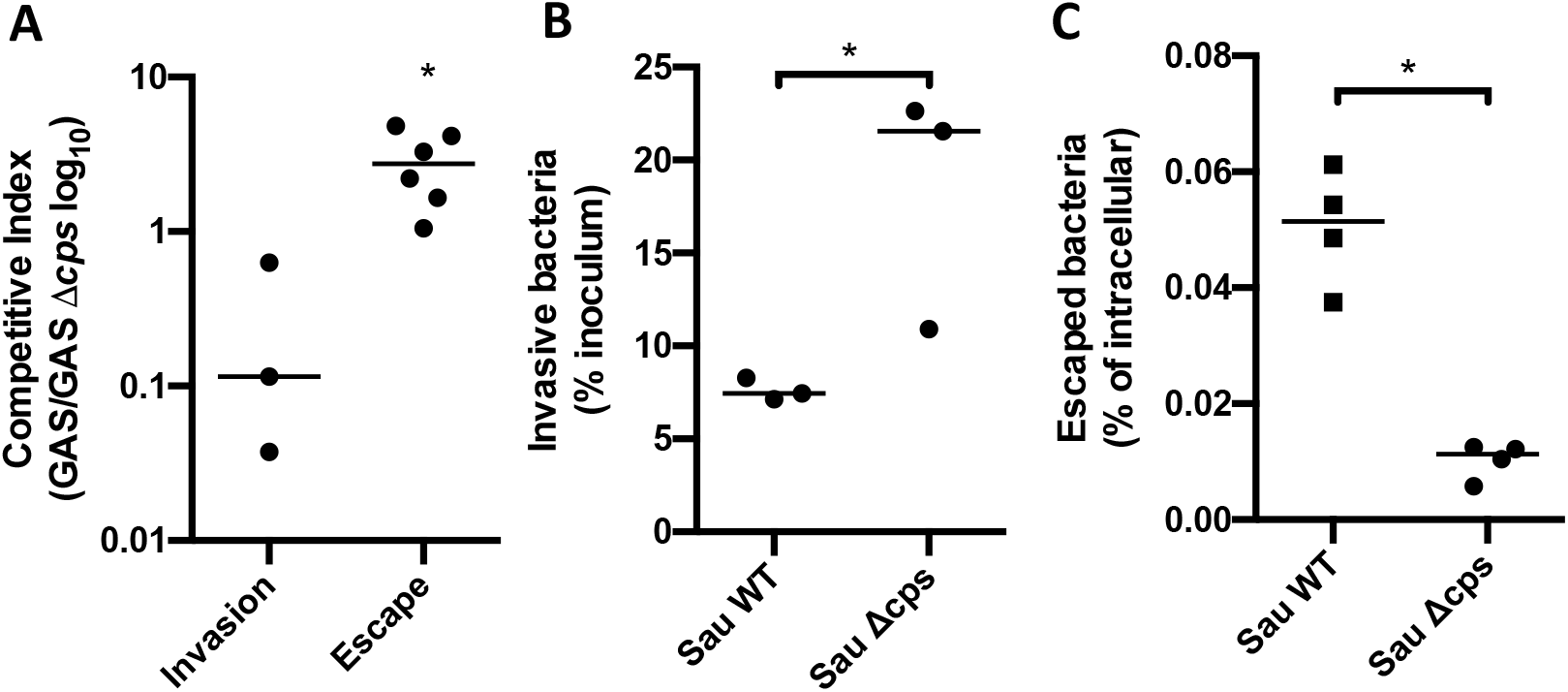
Capsule promotes transcytosis of *S. pyogenes* and *S. aureus*. **(A)** In competition (ratio 1:1), encapsulated *S.pyogenes* (GAS) escape at higher rates than a non-encapsulated isogenic mutant (GASΔcps), despite being initially outnumbered within the cells. **(B)** Encapsulated *S. aureus* are less internalized but, **(C)** escape at higher rate than non-encapsulated *S. aureus*. The use of unmarked mutants did not allow for competition assay. Statistical analyses: Mann-Whitney U-test. Errors bars represent standard error of the mean.

## DISCUSSION

Despite progress made with new vaccines and increasing access to intensive critical care, invasive bacterial diseases remain a leading cause of human morbidity and death worldwide (32, 33). Bacteremia, which can result in sepsis, also results in bacterial invasion of tissues (34). The latter alone can result in serious organ-specific complications. Examples include liver abscesses (35, 36), kidney damage and failure (37), cardiac microlesions and adverse cardiac events (23, 38), and meningitis (39, 40). Thus, it is imperative to improve our understanding of the molecular mechanisms underlying these processes in order to identify prophylactic strategies and prevent associated damage.

Studied for over a century, the capsule is one of *Spn*’s primary virulence determinants. The principal roles of the capsule are to resist entrapment in mucus during colonization and avoid killing by phagocytes (10). So far 100 distinct serotypes of *Spn* capsule have been identified, with serotype-specific resistance to complement deposition demonstrated to be one reason for why certain serotypes have a higher propensity to cause IPD (5). Other factors related with the propensity of the capsule to cause disease are its abundance and negative charge, both which impair its association with host cells. Importantly, and although required for survival within the bloodstream, other than the demonstration that capsule is generally inhibitory of bacterial adhesion, how capsule influenced *Spn* translocation across vascular endothelial cells during invasive disease was, up to this point, unknown.

Given capsule’s inhibitory effect on bacteria uptake by host cells, studies on *Spn* interactions with VEC have relied heavily on the use of non-encapsulated mutants (14, 16, 41–43). We suspect this is why the role of capsule as an intracellular antioxidant has not been previously described. A few reports did use encapsulated bacteria to study pneumococci translocation across VEC concluding to a meaningful impact. Fuchs et al. found that encapsulated *Spn* crossed human brain microvascular endothelial cell monolayers more efficiently than their unencapsulated counterparts (44). However, this study focused solely on serotype 7F, did not examine the mechanism, nor include findings with animal models. Ring et al. also described that different serotypes crossed an *in vitro* blood-brain barrier model with various efficiencies (22). However, their experiments emphasized the relevance of the genetic background showing that phase variation, known to modulate capsule and other surface determinants (18, 45), was a major factor in *Spn*’s ability to cause meningitis. Here, to overcome the genetic variability between strains with different capsule types, we used isogenic capsule switch mutants to study the contribution of capsule alone in the fate of *Spn* once within the cell.

Our experiments revealed that the capsule has a vital role during transcytosis through VEC layers by preventing, or at least delaying, *Spn* intracellular killing. Within VEC, *Spn* encounter a hostile environment with the presence of microbicidal factors such as free-radicals, antimicrobial peptides, lytic enzymes and low PH. Studies have shown that polysaccharides are efficient free radical scavengers (46, 47). Additionally, it has been described that capsule enlargement of the intracellular yeast, *Cryptococcus neoformans*, confers resistance to oxidative stress within the phagolysosome (48). Herein we show bacterial capsular polysaccharide also has this property and that this conferred intracellular protection within VEC, facilitating the bacterium’s translocation into organs from the bloodstream. Importantly, not all serotypes were equally capable of conferring protection against ROS. This result was expected given the considerable biochemical diversity of different serotypes. Along these lines, type 4 capsule was observed to confer superior antioxidant capabilities and resistance to intracellular killing versus type 2. The importance of this feature in pneumococcal pathogenesis was subsequently evidenced by our results in mice that showed enhanced cardiac invasion by type 4 carrying pneumococci and the fact that oxidative stress resistance and corresponding ability to translocate across VEC monolayers conferred by different serotypes in an isogenic background was positively correlated with the published attack rates in humans for the tested serotypes. Thus, this is a new and important feature of capsular polysaccharide that was not previously appreciated. Importantly, we demonstrated that capsule also promoted *S. pyogenes* and *S. aureus* escape from within MCEC cells, suggesting that our observation can be applied to other medically important bacteria that carry capsule.

The exact manner by which capsule conferred resistance to oxidative stress remains an open question and most varies considerably between serotypes. Nonetheless, we have gained some important insights. Foremost the amount of capsule produced matters. This was evidenced by our observation that reductions in capsule reduced the protective effects measured. Second, specific biochemical features of the capsule, such as acetylation, have relevant consequence and testing of other serotypes in an isogenic background that carry or lack these features is a possible path forward to determine their relevance. Third, capsule shedding promotes protection. The latter is consistent with similar studies that examined the protective effects of capsule against cationic antimicrobial peptides. Fourth, an exogenous source of capsule can protect unencapsulated *Spn* against ROS. While this may not be as critical during VEC translocation, this is likely to be an important trait within the nasopharynx and biofilms, where individual bacteria carry less capsule but are surrounded by the extracellular matrix that contains capsule (49). Ongoing efforts by our group are focused on the identification of the specific biochemical moieties in serotype 4 capsular polysaccharide that are altered as result of exposure to H_2_O_2_ – with the notion these are what confer protection. Moreover, testing if the presence of these specific residues in other serotypes are predictive of their antioxidant properties and ability to confer intracellular protection.

Capsules antioxidant effects were not absolute, and we did observe killing over time. This suggests that capsules role during VEC translocation is to delay killing versus promoting long-term intracellular occupancy. The former providing pneumococci with more time to complete the translocation process. Importantly, these antioxidant properties are likely distinct from other key features of the capsule – such as resistance to complement deposition and opsonophagocytosis. The *in vivo* importance of the latter evidenced by our results in mice that showed capsule serotype had strong impact on bacterial burden in the bloodstream following challenge. Thus, the biochemical properties of each serotype may favor or disfavor certain forms of disease due to how it influences interactions with host components and cells. Along such lines, our observations raise the question of the importance of capsule during the crossing of human epithelial barriers. Similar to what occurs in VEC, some capsule types may allow *Spn* to better access the bloodstream from the nasopharynx or the lungs through epithelial cell translocation. This is an important question to address as epithelial cell translocation is the first step towards invasive disease following respiratory tract infection.

Finally, the fact that serotype-conferred resistance to oxidative stress and intracellular survival *in vitro* is positively correlated with serotype attack rate in humans suggests that these traits can be used to our advantage. For example, we can screen and identify the non-vaccine serotypes which are most likely to be problematic in the future should serotype shift in response to the vaccine continue. Similar studies should be performed with other invasive pathogens such as group B streptococci (GBS), whose primary virulence determinant is also a capsular polysaccharide, thereby informing serotype inclusion in any future GBS vaccine as well. Testing for correlations between serotype-conferred resistance to oxidative stress and other features of the pathogenic process, such as the ability to overcome phagocytes, are now warranted for the same reasons.

In summary, we have identified a new and key role for pneumococcal capsule in promoting invasive disease. That is, capsular polysaccharide serves as an antioxidant and this facilitates bacterial translocation across VECs. Our finding on how specific biochemical attributes of capsule confer *Spn* protection may result in the identification of novel targets to block the development of bacterial invasive disease. Moreover, assist in the rationale design of future vaccines.

## MATERIAL AND METHODS

### Bacterial strains and growth conditions

Strains used in this study are described in **Supplemental Table 1**. Isogenic mutant derivatives of *Spn* serotype 4 strain TIGR4, were created using splicing overlap extension PCR as described and using primers listed in **Supplemental Table 2**. Isogenic capsule switches were constructed as described in supplemental material and methods. Bacteria were grown in Todd-Hewitt broth with 0.5% yeast extract (THY), or on blood agar plates (Remel), in a humidified atmosphere at 37°C with 5% CO_2_.When necessary, Chloramphenicol (4.5μg/ml), Erythromycin (0.5μg/ml), Kanamycin (200μg/ml), and Streptomycin (300μg/ml) were added to the media.

### Financial Support

Support for these experiments was provided through National Institutes of Health (NIH) grants AI114800, AI146149, AI148368 and AI156898 to CJO. Training of ANR is supported by NIH T32 AI007051. Training of KLK supported by NIH T32 HL134640.

### Statistical analyses

All analyses, excluding the NMR, were performed in GraphPad Prism 8 (San Diego, CA). Data plotted as mean of three technical replicates unless otherwise noted. Biological triplicates graphed with error bars denoting mean and standard error of the mean. Comparisons between two groups within one independent variable assessed by unpaired Student’s T-test when data were normally distributed or by Mann-Whitney U test when data distribution was not normal. Comparisons between three or more groups were assessed by ANOVA with Tukey’s post-test unless otherwise noted in the figure legend. Dependency between two variables were assessed by Spearman Correlation.

### Ethics statement

All mice experiments were reviewed and approved by the Institutional Animal Care and Use Committee at The University of Alabama at Birmingham, UAB (Protocol # IACUC-20175). Animal care and experimental protocols adhered to Public Law 89–544 (Animal Welfare Act) and its amendments, Public Health Services guidelines, and the Guide for the care and use of Laboratory Animals (U.S. Department of Health & Human Services).

### Mice experiments

Female 6 weeks-old C57B6 (Jackson) were challenged with 1.0×10^5^ pneumococci by intratracheal (IT) or intraperitoneal (IP) infections in 20μL or 100μL PBS respectively. Blood for assessment of bacterial burden was obtained by tail bleeds (XμL) every 12h. At final time point (T_0_+36h~42h) or when deemed moribund, 100μl of blood was collected via retro-orbital bleed from anesthetized mice for bacteremia determination. Mice were subsequently euthanized by CO_2_ asphyxiation and death confirmed by pneumothorax. Mice were perfused by cardiac puncture using PBS then collected organs were washed thoroughly with PBS and homogenized in 1mL of PBS for bacterial burden determination

### Cell culture

Mouse Cardiovascular Endothelial Cells (MCEC) were grown in DMEM (Corning) supplemented with 10% heat-inactivated fetal bovine serum (FBS; Atlanta Biologicals), and 1X penicillin/streptomycin solution (Corning, Cellgro) in a humidified atmosphere at 37°C and 5% CO_2_.

### Adhesion and invasion experiments

MCEC cells were seeded at 5.0×10^5^ cells per well in 12-well plates. Experiments were performed as described in (50) using 1.0×10^7^ Spn (MOI=10). Intracellular survival rates were determined normalizing the number of intracellular bacteria at any given timepoint normalized by the number of intracellular bacteria after 1 hour of incubation with bactericidal concentration of gentamicin.

### Transcytosis assays

5.0×10^5^ MCEC cells were seeded on Transwell^®^ permeable inserts (12mm, 3μm pore-size, Costar) in 12-wells and incubated for at least 48 hours at 37°C with 5%CO_2_. For each experiment, extra inserts were seeded to determine the number of intracellular bacteria. 5.0×10^6^ *Spn* were added to the cells before centrifugation at 500g for 5 minutes and incubation for 30 minutes at 37°C with 5%CO_2_. Inserts were then washed three times with prewarmed PBS and incubated for 30 more minutes in DMEM at 37°C with 5%CO_2_. Gentamicin was then added at a bactericidal concentration (200μg/ml) in the upper chamber and a bacteriostatic concentration (20μg/ml) in the lower chamber before incubation at 37°C with 5%CO_2_ for 1h. Filters were moved to new plates and incubated for 1 extra hour in DMEM containing bacteriostatic concentrations of gentamicin. Extra inserts seeded to determine the number of intracellular bacteria were washed three times in PBS after gentamicin incubation for 1 hour. Cells were lysed by addition of cold water and 15 minutes of incubation at 4°C before plating. The escape rate was determined by the number of CFU recovered in the lower chamber normalized by the number of intracellular bacteria.

### Stress tolerance assays

*Spn* from an overnight culture on a blood agar plate was used to inoculate THY and this preculture was incubated until OD620nm=0.3-0.4. For ROS assays, 5.0×10^7^ bacteria were mixed with 20mM H_2_O_2_ or menadione in THY (final concentration 10mM). Stress tolerance was determined by counting the number of viable bacteria at any given timepoint normalized by the number of bacteria incubated in plain THY.

### FITC-dextran exclusion assay

To quantify the capsule thickness, we measured the exclusion area of FITC-dextran (FD2000S, sigma) based on previous article (51). Briefly, *Spn* were cultured in THY media until OD600nm 0.3 and centrifuged at 3,000g for 10 min and pellet was resuspended in 500 μl of PBS or 4 % paraformaldehyde solution. 18 μl of resuspension was mixed with 2 μl FITC-dextran solution (10 mg/ml, final 1 mg/ml concentration). The mixed solution was put onto a microscope slide and visualized with Leica LMD6 microscope equipped with DFC3000G monochrome camera at 40x magnification. The obtained images were analyzed using ImageJ processing software.

### NBT reduction assay

To test the antioxidant capability of *Spn* capsule we used the NBT reduction assay coupled to NADH and PMS (52, 53). Briefly, the reaction was performed in 96-well plates using 200 μl per assay. A mix of NADH (166 μM) and NBT (43 μM) freshly prepared in phosphate buffer (40 mM, pH 7.6) was incubated for 2 minutes at room temperature and NBT reduction was started by the addition of 2.7 μM PMS. The plates were read in an iMark Microplate Reader (Bio-Rad) at 37°C. Optical density was monitored at 550 nm every 30 seconds for 30 minutes. The antioxidant efficacy of the purified capsule was estimated by the capability of protection NBT from reduction compared to the controls with no polysaccharide.

### NMR Spectroscopy

Control sample of type 4 capsule polysaccharide (ATCC, Manassas, VA) was prepared by dissolving ~6 mg in 3 mL of milli-Q water and dialyzed (Dialysis tubing: 8,000 molecular weight cut-off) overnight in 4L milli-Q water. Sample was then lyophilized and dissolved in 0.5 mL of 99.99% D2O (Cambridge Isotope Laboratories). For polysaccharide oxidation, ~6 mg of type 4 capsule polysaccharide was treated with 10 mM H_2_O_2_ (Alfa Aesar, Massachusetts) in a total 3 mL volume. Solution was incubated at 37 °C for either 30 mins or 3 hours. Solution was then dialyzed overnight against 4L of milli-Q H_2_O followed by lyophilization. Sample was then dissolved in 0.5 mL of 99.99% D2O. ^1^H-^1^H and ^1^H-^13^C NMR data were collected at 50 °C on a Bruker Avance II (700 MHz ^1^H) spectrometer equipped with cryogenic triple-resonance probes, processed with NMRPIPE (54) (and analyzed with NMRVIEW (55). HDO signal was used as a reference.

### Capsule polysaccharide purification

Capsule PS was purified from BLS101 (serotype 33A) and BLS104 (33X2, a variant of serotype 33A with defective wcjE and wciG) as described previously (29). Each strain was grown in 2 liters of a chemically defined medium (56), supplemented with choline chloride (1 g/L), sodium bicarbonate (2.5 g/L), and cysteine HCl (0.73 g/L). After overnight incubation, the bacteria were separated from the supernatant by centrifugation (15,344 × *g*; 30 min; 4°C), resuspended in 0.9% NaCl with sodium deoxycholate (0.05%) and mutanolysin (100 U/ml), and incubated at 37°C for 72 h. The lysate was centrifuged, dialyzed, and subjected to ion-exchange chromatography. Since serotype 33A and 33X2 capsules are neutral polysaccharides, they did not bind to the DEAE column and were recovered in flowthrough and wash fractions. Fractions containing 33A PS were detected and quantified with the serotype 33A inhibition ELISA assay using factor sera 20b as the serological marker. Whereas, 33X2 PS containing fractions were detected by anthrone assay (57). To ensure purity, the fractions were tested for the presence of cell wall polysaccharide (CWPS) by an inhibition ELISA assay using phosphocholine specific monoclonal antibody (HPCG2b). The fractions with high polysaccharide concentration, low CWPS concentration, and low absorbance at 260 and 280 nm were pooled, dialyzed, and lyophilized.

### Study approval

This study was carried out in accordance with the recommendations of the University of Alabama at Birmingham Institutional Animal Use and Care Committee (IACUC) in compliance with the federal regulations set forth in the Animal Welfare Act and the recommendations of the National Institutes of Health Guide for the Care and Use of Laboratory Animals. All procedures used in this study were approved by the IACUC under protocols 21850 and 20175.

## Author contribution

TB, EM, KLK, ANR, FG, HI, SB, NMA, and BLS carried out the experiments. TB, EM, JSS, MHN and CJO and contributed to the design and conceptualization of the project. TB, EM and CJO wrote the manuscript. All authors provided critical commentary to help shape the manuscript.

## SUPPLEMENTAL FIGURE LENGENDS

**Figure S1.**
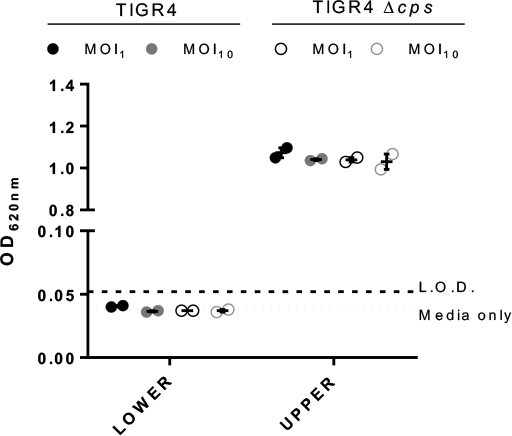
Representative transcytosis post assay experiment. 0.5% Evans blue was added to the upper Transwell chamber and incubated for 1h at 37°C. OD_620nm_ of media from the lower layer was measured and compared to a reference Evans blue standard.

**Figure S2.**
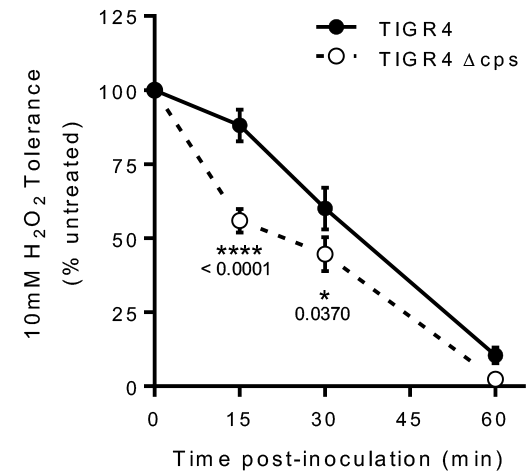
Reactive oxygen species tolerance was determined by calculating the number of live *Spn* in THY supplemented with H_2_O_2_. Errors bars represent standard error of the mean. Statistical analyses: 2way ANOVA with repeated measures

**Figure S3.**
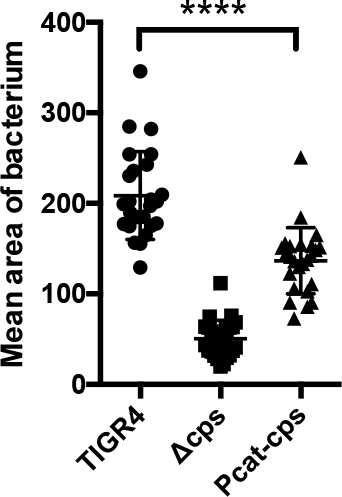
FITC-dextran exclusion confirmed production of a reduced capsule amount of TIGR4 expressing cps locus under the control of Pcat promotor (Pcat-*cps*). Each point represents an individual bacterium. Statistical analyses: Mann-Whitney U-test.

**Figure S4.**
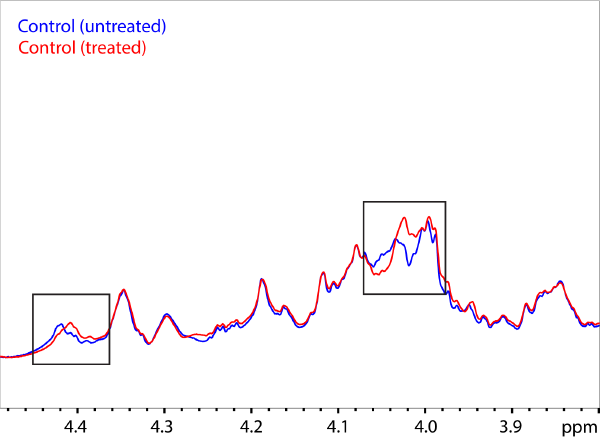
NMR analysis of type 4 capsule treated with 10mM H_2_O_2_ for 30 minutes versus control.Differences are boxed.

**Supplemental Table 1.**
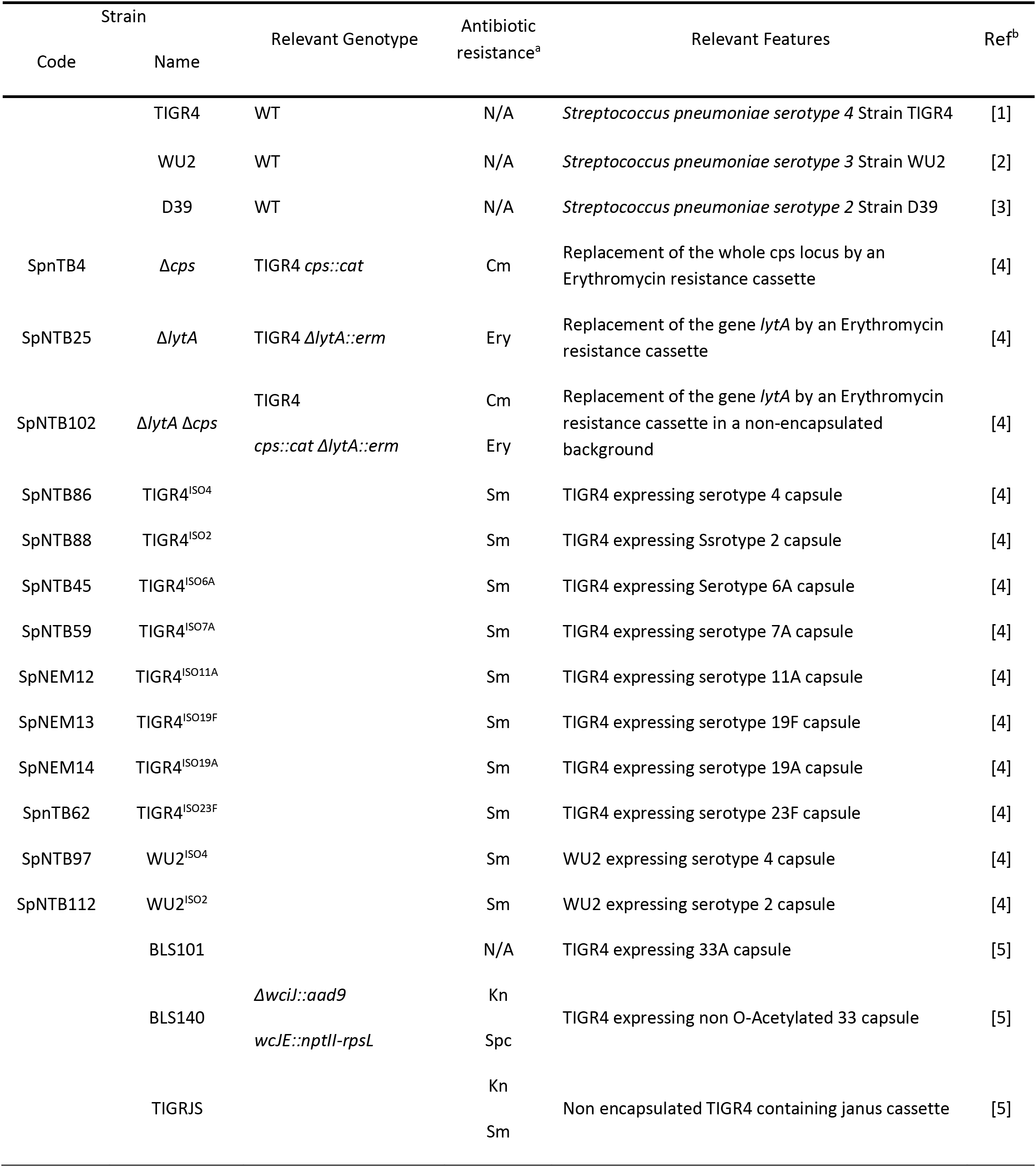
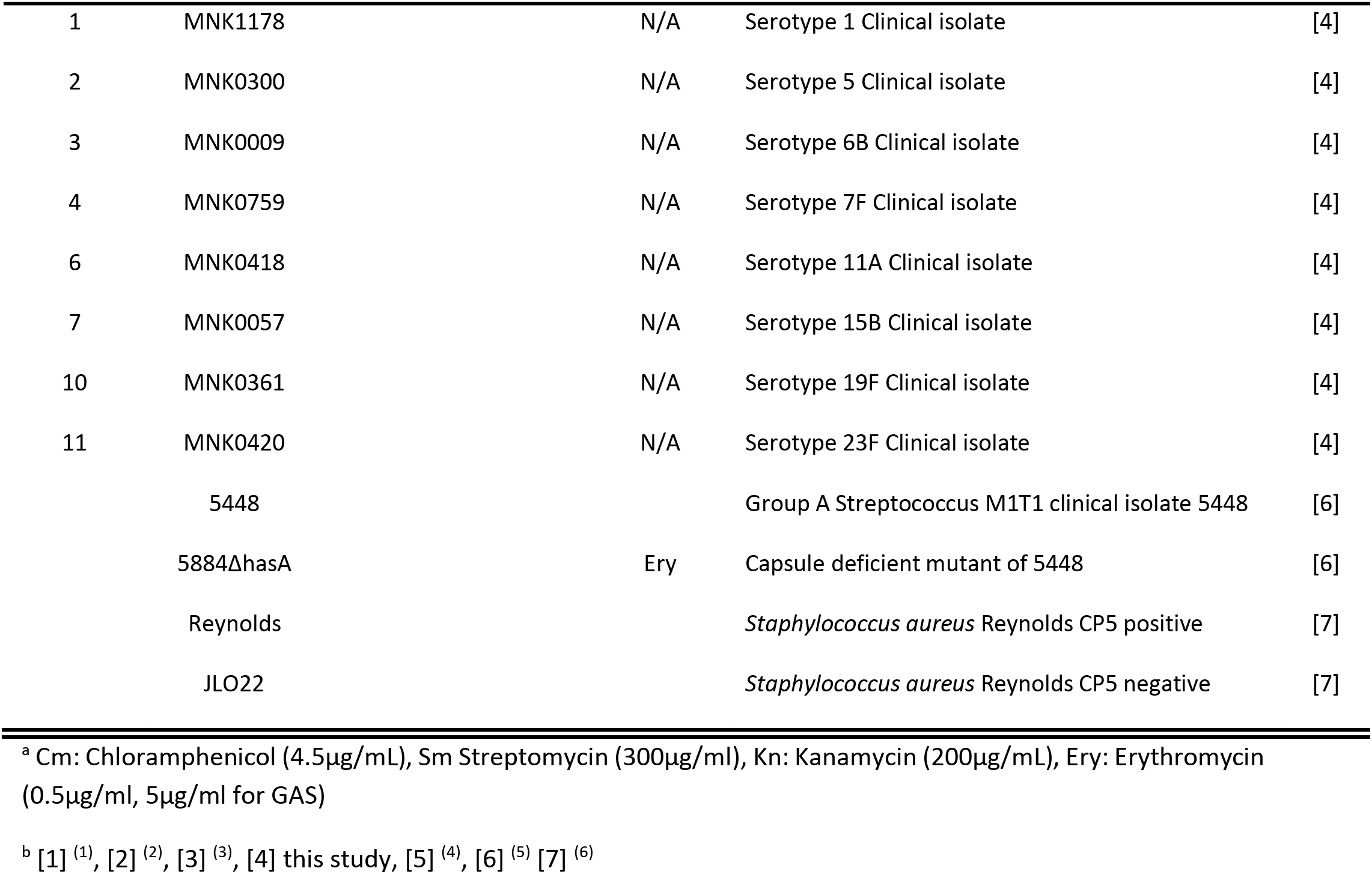
Strains used in this study.

---

**Supplemental Table 2.**
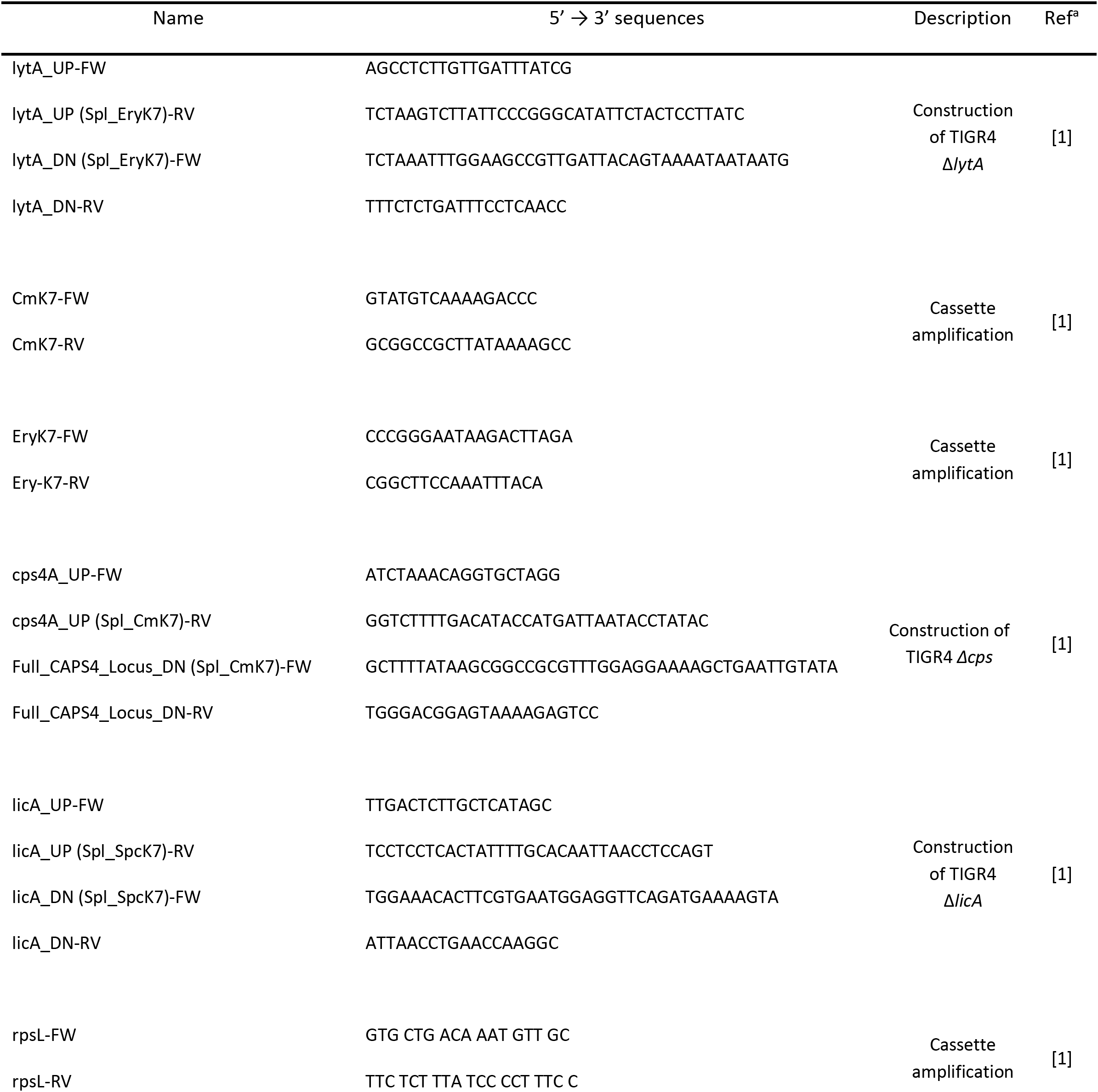
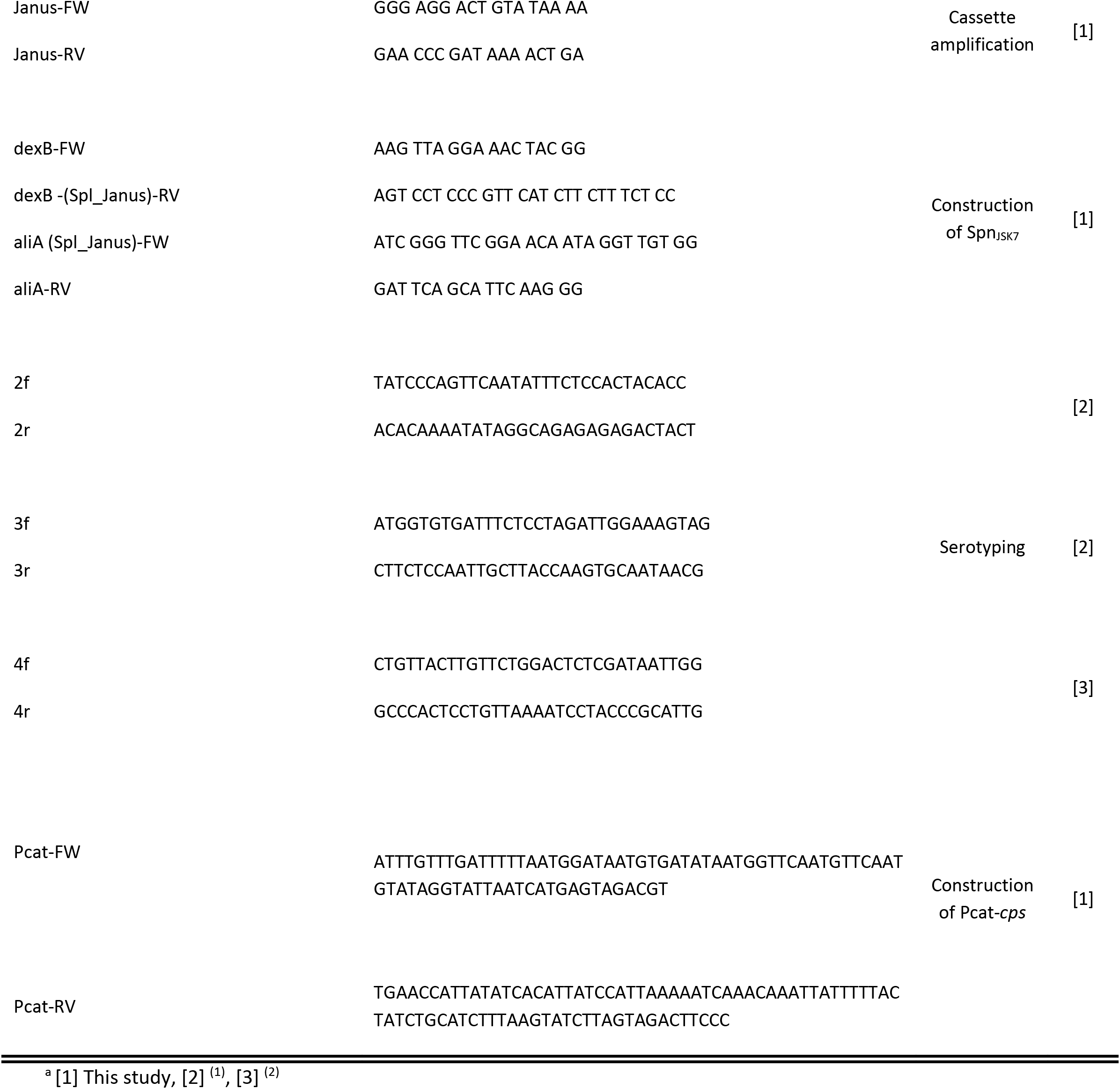
Primers used in this study.

